# Tissue-specific immune transcriptional signatures in the bordering tissues of the mouse brain and retina

**DOI:** 10.1101/2024.06.30.601438

**Authors:** Fazeleh Etebar, Paul Whatmore, Damien G. Harkin, Samantha J. Dando

## Abstract

**Background:** Bordering the central nervous system (CNS) parenchyma are the pia mater (the innermost layer of the meninges enveloping the brain) and the choroid (underlying the retina). While near the neural parenchyma, the pia mater and choroid are external to the immune privileged environment of the brain and retina and thus are distinct immune compartments. This study aimed to characterise the transcriptomic signatures of immune cells within the pia mater and choroid bordering the healthy adult mouse CNS.

**Methods:** Brains and eyes were obtained from 7-week-old female C57Bl/6J mice. Pia mater-enriched tissue and choroid were dissected and processed for fluorescence activated cell sorting of CD45^+^ immune cells and single cell RNA-sequencing. Additionally, single cell RNA-sequencing was performed on immune cells isolated from choroid obtained from human donor eye tissue. Immunostaining and confocal microscopy of wholemount tissue were used to validate selected immune cell populations *in situ*.

**Results:** A total of 3,606 cells were sequenced from mouse tissues, including 1,481 CD45^+^ cells from pia mater-enriched tissue and 2,125 CD45^+^ cells from choroid. Clustering and differential gene expression analysis revealed heterogeneous subtypes of monocytes/macrophages, dendritic cells, T cells and B cells. While some clusters were common to both pia mater and choroid, others exhibited tissue-specific gene expression profiles and potential functional specialisations. Analysis of 6,501 CD45+ cells sequenced from human choroid identified similar immune cell populations to mouse choroid.

**Conclusions:** This study provides a detailed characterisation of the molecular signatures of immune cells within the vascular connective tissues bordering the healthy brain and retina, and their potential roles in immune protection.

## BACKGROUND

The central nervous system (CNS; comprising the brain, spinal cord and neural retina) is a complex organ system that requires specialized support from bordering tissues and spaces. The bordering tissues of the brain include the meninges, comprising three layers that cover the brain; and the choroid plexus within the ventricles, which secrete cerebrospinal fluid (CSF) and form the blood-CSF barrier (1, 2). Bordering the retina is the choroid, a highly vascularised layer underlying the retinal pigment epithelium; and the ciliary body, which secretes aqueous humour and forms part of the blood-ocular barrier (3, 4). The pia mater and arachnoid mater (the leptomeninges) are the inner and middle layers of the meninges respectively. Both the leptomeninges of the brain and choroid of the eye have vastly different immune properties compared to the underlying neural parenchyma (4). As the immunological interface between the periphery and neural environment, the leptomeninges and choroid play important roles in immune surveillance and defence against pathogens and insults (3, 5).

Beyond their role in immune defence, tissue resident immune cells contribute to homeostatic functions (6, 7). Defining the diversity of immune cells within the meninges and choroid is therefore important for advancing our understanding of the immunobiology at the borders of the brain and retina. Unlike the CNS parenchyma, which is populated by microglia, the meninges and choroid contain macrophages, dendritic cells, mast cells and lymphocytes (7–11). Seminal studies of border-associated macrophages (BAMs) within the mouse meninges, choroid plexus and perivascular space demonstrated that these cells exhibit region-specific transcriptomic signatures and ontogenies (8, 9). However, comparative studies of immune cells within the bordering tissues of the brain and retina are lacking. Since increasing evidence suggests that immune cell identities are shaped by their tissue environment, we hypothesised that the leptomeninges and choroid would contain similar types of immune cells but would be distinguished by tissue-specific gene expression profiles.

Advances in single-cell RNA sequencing (scRNA-seq) technology have enabled more detailed molecular characterisation of immune cell subtypes at an unprecedented level of resolution. We therefore used scRNA-seq to profile the immune landscape of the pia mater-enriched leptomeninges and choroid bordering the healthy mouse brain and retina, and identify tissue-specific transcriptomic signatures.

## METHODOLOGY

### Mice, tissue collection and dissection

Mouse experiments were approved and performed in accordance with the QIMR Berghofer Medical Research Institute Animal Research Ethics Committee (approval A18613M); QUT AEC administrative approval 1800001261. A notifiable low risk dealing for the use of genetically modified mice in this project was obtained from the QUT University Biosafety Committee (NLRD approval 1800000957). Adult (7 weeks of age) female C57Bl/6J mice were used for this study. C57Bl/6J mice were purchased from the Animal Resources Centre (Canning Vale, Western Australia). Mice were maintained on a 12:12 hour light cycle with access to food and water *ad libitum*. For experiments, mice were deeply anaesthetised by an intraperitoneal injection of sodium pentobarbital (150 mg/kg). Mice were then perfused immediately through the left ventricle with cold phosphate buffered saline (PBS) to clear the vasculature of blood components including circulating leukocytes. When tissues were being collected for histology and immunostaining, mice were also transcardially perfused with 4% paraformaldehyde (PFA) to deliver fixative systemically to the animal.

Brains and eyes were collected from perfused mice and dissected under a stereomicroscope. During mouse brain collection, the calvaria (containing dura mater and likely part/all of the arachnoid) (12, 13) was removed, leaving predominantly the pial leptomeninges adherent to the surface of the brain. To obtain samples of pia mater-enriched tissue, the superior surface of the cerebral cortex was trimmed using a flat-edged razor blade as previously described (14). The resulting sample that was processed for scRNA-seq was enriched for pia mater but also contained a small amount of underlying cortical tissue. Choroids were collected as previously described (15).

### scRNAseq of immune cells within the pia and choroid

Immune cells (CD45^+^ cells) were isolated from the choroid and pia mater-enriched tissue obtained from n=20 C57Bl/6J female mice by FACS. Freshly dissected choroid and pia mater-enriched tissue were pooled to obtain a sufficient number of cells. The dissected tissues were transferred to 50 mL tubes containing dissection buffer (1x HBSS, no calcium, no magnesium, no phenol red; 5 mM glucose; 15 mM HEPES) and all processing steps were performed on ice. The tissues were passed through a 70 μm nylon cell strainer (Falcon 352350) using the plunger of a 5 mL syringe to obtain single cell suspensions, and then pelleted by centrifugation at 400 *g* for 5 min at 4 °C. Pia mater-enriched tissue was subsequently resuspended in 30% (v/v) Percoll (GE Healthcare 17089102) in 1 x PBS and centrifuged at 700 *g* for 10 min at 4 °C without a brake being applied. The small, myelinated layer formed at the top of the tube (from cortical tissue) was discarded prior to resuspending the cell pellet.

The number of viable cells was determined by exclusion of Trypan blue stain as viewed by microscopy, assisted by use of a hemocytometer. Cells were subsequently resuspended in FACS buffer (3 mM EDTA, 0.1% (w/v) BSA, 1 x PBS, 100 μg/mL DNase I) and stained with rat anti-mouse CD16/CD32 (BD Biosciences 553141) for 15 mins, then centrifuged at 400 *g* for 5 mins. The pellet was resuspended in FACS buffer containing BV421 rat anti-mouse CD45 antibodies (BD Biosciences 563890), incubated for 30 min on ice, then centrifuged and resuspended in FACS buffer. CD45^+^ were isolated from stained cell suspensions using a BD FACS Aria IIIu (100 µm nozzle) at QIMR Berghofer Flow Cytometry Facility. Dead cells were identified and excluded during sorting by staining with propidium iodide (1 μg/ mL, BD Biosciences 556463).

Sorted cells were partitioned into single cell droplets using the 10X Genomics Chromium Controller. Subsequently, scRNA-seq libraries were prepared using a Chromium single cell reaction 3’ v3.1 kit (10X Genomics). Single cell libraries were sequenced on an Illumina NovaSeq S1 instrument, producing paired-end reads.

### scRNA-seq of human choroidal immune cells

Human tissue experiments were approved by Metro South Human Research Ethics Committee (HREC/07/QPAH/048); QUT HREC administrative approval 0800000807/ERM Project No. 5297. For study of human choroidal immune cells, scRNA-seq was performed on cells isolated from human post-mortem eyes. Fresh (unfixed) human eye cups from a 45-year-old male donor with no history of eye disease or diabetes was obtained within 17 hours post-mortem from the Queensland Eye Bank. The vitreous and retina were removed from the eye cups and choroid-RPE was subsequently dissected from the sclera. Choroid-RPE tissue was transferred into a 12-well plate and subjected to enzymatic digestion with collagenase type II (200 U/mL in HBSS (calcium, magnesium, no phenol red)) for 60 minutes at 37 °C, with tissue triturated every 15 minutes. After 60 minutes, choroidal tissue was triturated with a pipette and then passed through a 70 µm cell strainer to generate a single cell suspension. The cell suspension was pelleted by centrifugation at 400 *g* for 5 mins, resuspended in freezing medium (DMEM/F12 containing 10% (v/v) DMSO and 10% (w/v) bovine serum albumin) and transferred to a Cool Cell freezing chamber at -80 °C for controlled freezing. Cells then were transferred to liquid nitrogen storage. For scRNA-seq studies, cells were thawed and pre-warmed DMEM/F12 containing 10% (w/v) BSA was added in a dropwise manner. The viability and yield of thawed cells were determined by Trypan blue staining using a haemocytometer.

Human choroidal immune cells were isolated from thawed cell suspensions by FACS. Thawed cell suspensions were incubated with anti-human Fc Block (BD Biosciences 564220) for 15 mins, then centrifuged at 400 *g* for 5 mins. The pellet was resuspended in FACS buffer containing BV421 mouse anti-human CD45 antibodies (BD Biosciences 563879) and incubated on ice for 30 mins. Cells were centrifuged at 400 *g* for 5 mins then resuspended in FACS buffer for sorting. Live (propidium iodide negative) immune cells (CD45^+^) were sorted using a BD FACS Aria IIIu with a 100 µm nozzle.

The isolated cells were partitioned into single cell droplets using the 10X Genomics Chromium Controller and scRNAseq library preparation was completed using the Chromium 3’ v3.1 single cell reaction kit (10X Genomics). The single cell library was sequenced on an Illumina NovaSeq S1 instrument, producing paired end reads.

### scRNAseq data analysis: Processing, QC metrics, removing non-target cells and clustering

Sequences were initially processed using 10X Genomics Cell Ranger v7.0.0 software (16) on a Linux HPC (high performance computing) environment. The Cell Ranger workflow consisted of two primary steps: first ‘cellranger mkfastq’ was used to demultiplex raw base call data files and convert these to fastq format files; second, ‘cellranger count’ was used to align sequences to the mouse reference genome, and then quantify the number of aligned sequences per genomic feature (i.e. defined gene regions), producing a count table that formed the basis of further downstream analysis (dimensionality reduction, clustering, and visualization). Additionally, ‘cellranger aggr’ was run to aggregate the output from multiple cellranger count runs (samples). Counts were re-quantified based on relative library size (normalisation) and gene expression was re-calculated.

The count table generated by Cell Ranger was imported into R v4.0.5 (R Core Team, 2022) (17) using the ‘Read10X’ function from the Seurat package (18). This was combined with the cell IDs (barcodes.tsv.gz) the gene IDs (features.tsv.gz) into a single Seurat database object using the ‘CreateSeuratObject’ function. Raw count data were transformed via dimensionality reduction so Principal Component Analysis (PCA), t distributed stochastic neighbour embedding (t-SNE) and Uniform Manifold Approximation Projection (UMAP) plots could be generated. This involved: (1) normalisation of data by log transformation, (2) identification of genes that exhibited high cell-to-cell variation, (3) scaling the data so that highly expressed genes didn’t dominate the visual representation of expression, (4) performing the linear dimensional reduction that converts expression to dimensions, and (5) plotting the first two dimensions in a PCA, t-SNE or UMAP plot. Each of these steps was achieved using Seurat functions.

Prior to clustering, filtration was applied to identify and remove outlier cells with aberrantly high (> 4000) or low (< 200) gene counts. Optimal clustering was guided by the Clustree package (19), which facilitated identification of optimal resolution scores to prevent over-or under-clustering for each dataset. Once quality filtration was completed, cells were clustered by gene expression using Seurat’s ‘FindClusters’ function and clustering was visualised with t-SNE, dot plots and heat maps.

### scRNAseq data analysis: Identifying DEGs, cell subtypes, and functional enrichment

Differentially expressed genes (DEGs) were identified using Seurat’s ‘FindMarkers’ function. Significantly differentially expressed genes were defined as having an adjusted p < 0.05 and log FC > 0.5. Initially, clusters were identified as immune cell types based on their expression of common signature genes (Supplementary Table 1).

DEGs were then examined for functional enrichment in Kyoto Encyclopedia of Genes and Genomes (KEGG) pathways and Gene Ontology (GO) terms using the package clusterProfiler (20). Annotated KEGG pathway maps were generated using the package pathview (21). Annotation of primary gene IDs (gene symbol) to other gene identifiers (such as Entrez gene ID, Ensembl gene ID) was completed using the AnnotationHub package (22).

### Spatial validation of scRNA-seq data using immunohistochemistry and confocal microscopy

Immunohistochemistry was performed on fixed wholemount choroid and pia mater-enriched tissue obtained from C57Bl/6J mice. CD19 polyclonal antibodies (Invitrogen PA5-114970) and CD3 monoclonal antibodies (Invitrogen 14-0032-82) were used to label B cells and T cells, respectively. To achieve this, wholemount choroid and pia mater-enriched tissue were permeabilised in 0.5% (v/v) Triton x-100 (Sigma Aldrich X100-100ML) in PBS at room temperature for 1 h. The tissues were then blocked in 3.0% (w/v) bovine serum albumin (Sigma Aldrich A7906-500G) and 0.3% (v/v) Triton X-100 in PBS for 1 h at room temperature. Tissues were then incubated with primary antibodies overnight at 4 °C. Tissues were washed three times (each 10 min) in PBS, and then incubated with fluorophore-labelled secondary antibodies Goat anti-rabbit AF Plus 488 (Life Technologies A32731) for CD19, and Goat anti-rat AF647 (Life Technologies A21247) for CD3, along with Hoechst (20mM, Life Technologies 62249) for 2 h at room temperature. Tissues were washed three times (each 10 minutes) in PBS, mounted on microscope slides (SuperFrost Plus White 1mm slide, Thermo Fisher Scientific MENSF41296SP) and coverslipped (Coverglass, rectangular, 22 x 40 mm, 1.5 thickness, ProSciTech G425-2240). Samples were imaged using an inverted SP5 5-channel confocal microscope (Leica Microsystems). Z stacks were captured every 1 µm, with a line averaging value of 3 applied. Maximum intensity projection images were created using FIJI (23).

### Data availability

The scRNA-seq data generated in this study have been deposited in NCBI’s Gene Expression Omnibus (24) and are accessible through GEO Series accession number GSE253419 (https://www.ncbi.nlm.nih.gov/geo/query/acc.cgi?acc=GSE253419).

## RESULTS

### Identification of immune cell types in mouse pia mater-enriched tissue and choroid

To characterise immune cell populations in the bordering tissues of the mouse brain and retina, we sorted CD45^+^ immune cells from pia mater-enriched brain tissue and the choroid of 7-week-old female C57Bl/6J mice and performed scRNA-seq. Unsupervised clustering and *t*-distributed stochastic neighbour-embedding (tSNE) projections were performed on aggregated data from both tissues (3,606 total cells; Figure 1a), and clusters were identified based upon their expression of known marker genes (Supplementary Table 1). Significant overlap was observed between immune cell types isolated from pia mater-enriched tissue and choroid; however, some tissue-specific differences were detected (Figure 1b). Both tissues contained populations of monocytes/macrophages (MC/Mφ), dendritic cells (DCs), T cells, B cells, and natural killer cells (NK), whereas neutrophils were solely detected within pia mater-enriched tissue. Small numbers of mast cells were detected in both tissues (mostly in choroid); however, these did not form distinct clusters within tSNE plots. Microglia (MG) from the underlying neural tissue were also detected, particularly in pia mater-enriched tissue, which cannot be completely peeled away from the surface of the mouse brain.

**Figure 1.**
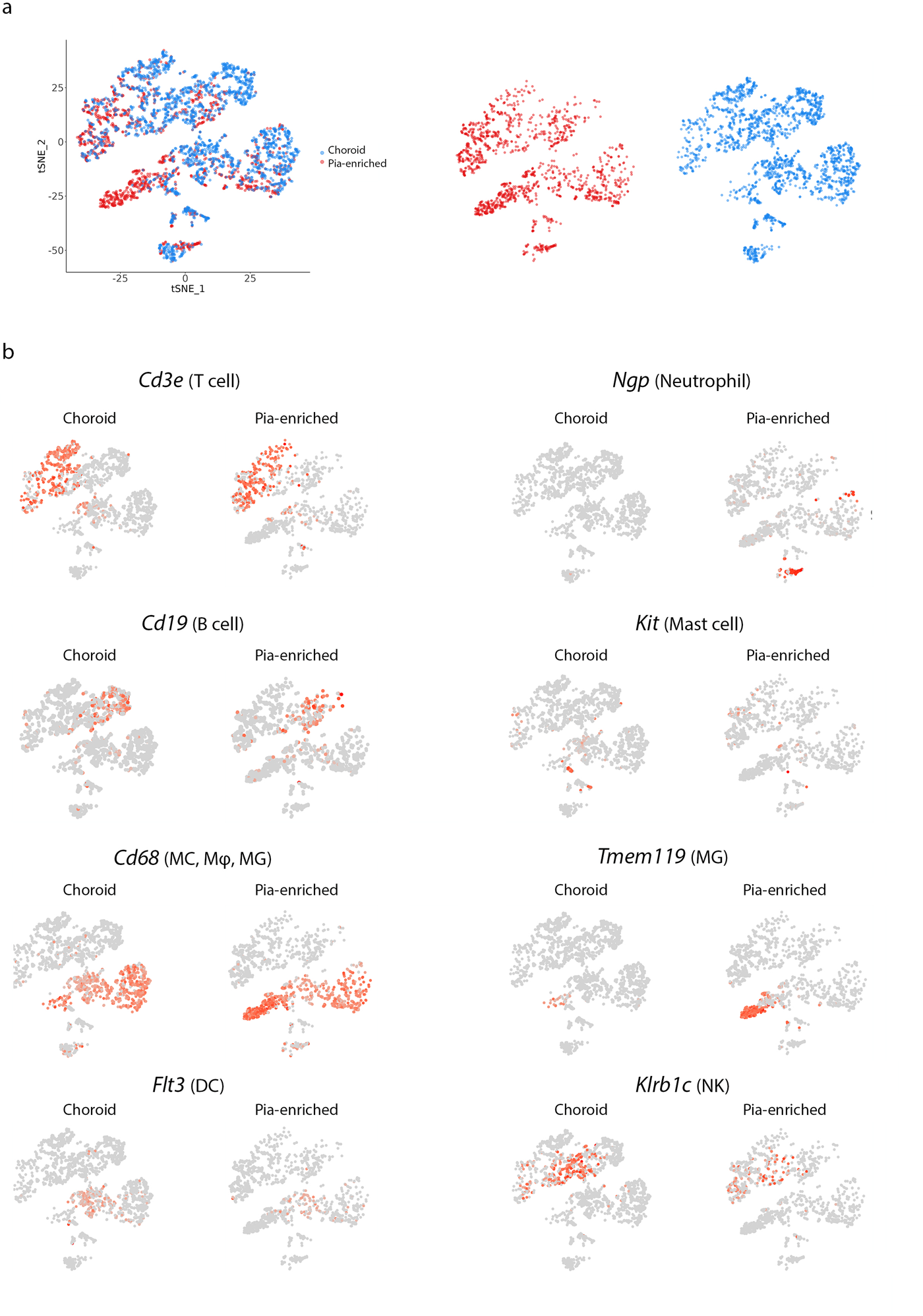
Comparison of immune cell types in mouse pia mater-enriched tissue and choroid. **(a)** tSNE plot of 3,606 immune cells (CD45^+^) sorted from pia mater-enriched brain tissue and the choroid of C57BL/6J mice (tissue pooled from n = 20 mice). Cells are coloured according to their origin (choroid = blue, pia mater-enriched tissue = red). **(b)** tSNE maps showing the frequency of cells expressing genes used to identify immune cell clusters (grey dots indicate cells not expressing the gene; red dots indicate cells expressing the gene). Monocyte, MC; Macrophages, Mφ; Dendritic cells, DCs; Natural killer cells, NK; Microglia, MG.

### Characterisation of immune cell clusters in mouse pia mater-enriched tissue and their unique transcriptomic signatures

To generate a transcriptomic profile of leukocytes within the homeostatic adult mouse pia mater, cells from pia mater-enriched tissue were re-analysed in a single dataset separate from choroidal leukocytes. Unsupervised clustering and tSNE projections were performed on 1,481 CD45^+^ cells (324,070 mean reads per cell) (Figure 2a), and clusters were identified based upon the expression of cell-specific genes (Figure 2b-e). Eleven immune cell clusters were detected including T cells (*Cd3e*^+^), B cells (*Cd79a*^+^ and *Cd19*^+^), MC/Mφ (*Cd14*^+^ and *Cd68*^+^), DCs (*Flt3*^+^), neutrophils (*Ngp*^+^) and NK cells (*Prf1*^+^ and *Klrb1c*^+^). Several subtypes of myeloid cells and T cells were further resolved based on differentially expressed genes (DEGs).

**Figure 2.**
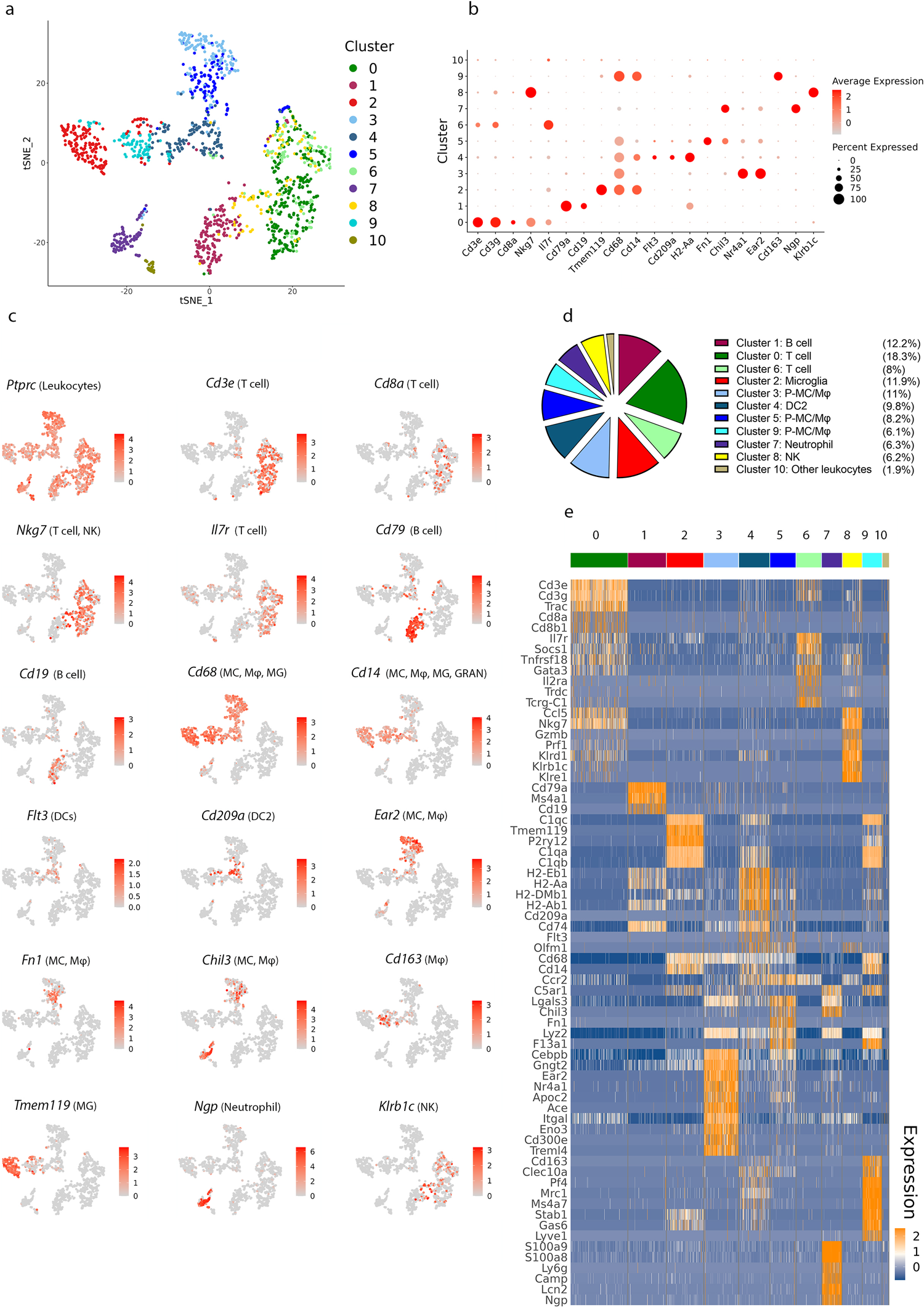
Identifying immune cell types in mouse pia mater-enriched tissue based on differentially expressed genes. **(a)** tSNE plot of 1,481 immune cells (CD45^+^) sorted from pia mater-enriched brain tissue pooled from n = 20 C57Bl/6J mice, showing the eleven immune cell clusters that were identified via unsupervised clustering. **(b)** Dot plots demonstrating the expression of cell type-specific genes, with the dot size representing the percentage of cells expressing the gene and the colour representing its average expression within a cluster. **(c)** tSNE maps showing the expression of key marker genes for the immune cell populations that were identified in (b). Grey, low expression; red, high expression, and range of log_2_ normalized counts are shown on the right of each plot. **(d)** Pie chart showing the proportions of different immune cell clusters within pia mater-enriched tissue. Clusters were identified based on expression of specific genes (the pie chart legend is ordered according to cell type rather than by numerical order of clusters). The frequency of each cluster is shown as percentage of the total immune cell population to the right of each cluster name. Note that the cluster identified as ‘other leukocytes’ expressed the CD45 gene (*Ptprc*), but none of the other examined lineage markers. **(e)** Heatmap of normalized expression for selected genes in each cluster. Range of log_2_ normalized counts are shown at right hand of heatmap (blue, low expression; orange, high expression). Monocyte, MC; Macrophages, Mφ; Dendritic cells, DCs; Natural killer cells, NK; Microglia, MG; granulocytes, GRAN.

Clusters 3, 4, 5 and 9 were identified as myeloid cells, which were the most abundant immune cells detected within pia-enriched tissue. Pial cluster 4 expressed *Flt3*, which is required for DC development and homeostasis (25), and had a transcriptional profile consistent with conventional type 2 DCs (cDC2s). This cluster was distinguished by 133 DEGs including *Cd209a*, *Cd74*, and antigen processing and presentation genes *H2-Eb1*, *H2-Aa*, *H2-Ab1*, and *H2-DMb1*. Three distinct pial MC/Mφ clusters were identified (cluster 3, 5, 9); each of these clusters expressed the common MC/Mφ gene *Cd68* and had low expression of *Flt3*. Pial MC/Mφ cluster 3 expressed 678 DEGs, including *Ear2* and *Nr4a1*, and other genes including *Gngt2*, *Apoc2*, *Ace*, *Itgal*, *Eno3*, *Cd300e* and *Treml4*. Pial MC/Mφ cluster 5 was characterised by 404 DEGs including *Chil3* and *Fn1*. Finally, pial MC/Mφ cluster 9 was distinguished from other pial MC/Mφ s by expression of complement C1q genes *C1qa*, *C1qb* and *C1qc*. This cluster featured 883 DEGs, which also included *Cd163*, *Clec10a*, *Pf4*, *Mrc1*, *Ms4a7*, *Stab1*, *Gas6*, and *Lyve1*.

Two pial clusters (cluster 0 and cluster 6) were identified as T cells based on their expression of *Cd3e*, *Cd3g* and *Cd3d*. Cluster 0 expressed genes consistent with CD8^+^ T cells (*Cd8a*, *Cd8b1*, and *Nkg7*), and effector memory T cells (*Il7r*), and was enriched for *Ccl5* expression. In contrast, cluster 6 T cells were characterised by expression of *Socs1*, *Tnfrsf18*, *Il2ra* (enriched in activated CD4^+^ T cells and CD4^+^ regulatory T cells) and *TcrgC1* (the gamma T cell receptor gene that is expressed by gamma-delta T cells). These findings suggest that the two identified pial T cell clusters contain various smaller subtypes of T cells.

### Characterisation of immune cell clusters in the mouse choroid and their unique transcriptomic signatures

Using the same approach as described above for pia-enriched tissue, a transcriptomic profile of leukocytes within the homeostatic mouse choroid was generated by performing unsupervised clustering and tSNE projections on 2,125 CD45^+^ cells (229,696 mean reads per cell) (Figure 3a). Thirteen immune cell clusters were resolved, and these clusters were identified based on their expression of cell-specific genes, including various subtypes of B cells (*Cd79a*^+^ and *Cd19*^+^), T cells (*Cd3e*^+^), DCs (*Flt3*^+^), MC/Mφ/ granulocytes (GRAN) (*CD68*^+^ and *Cd14*^+^), NK cells (*Klrb1c*^+^ and *Gzmb*^+^), and mast cells (*Kit*^+^) (Figure 3b-e). Similar to pia mater-enriched tissue, myeloid cells and T cells were the most abundant immune cell types within the choroid (Figure 3d) and corresponded to several clusters.

**Figure 3.**
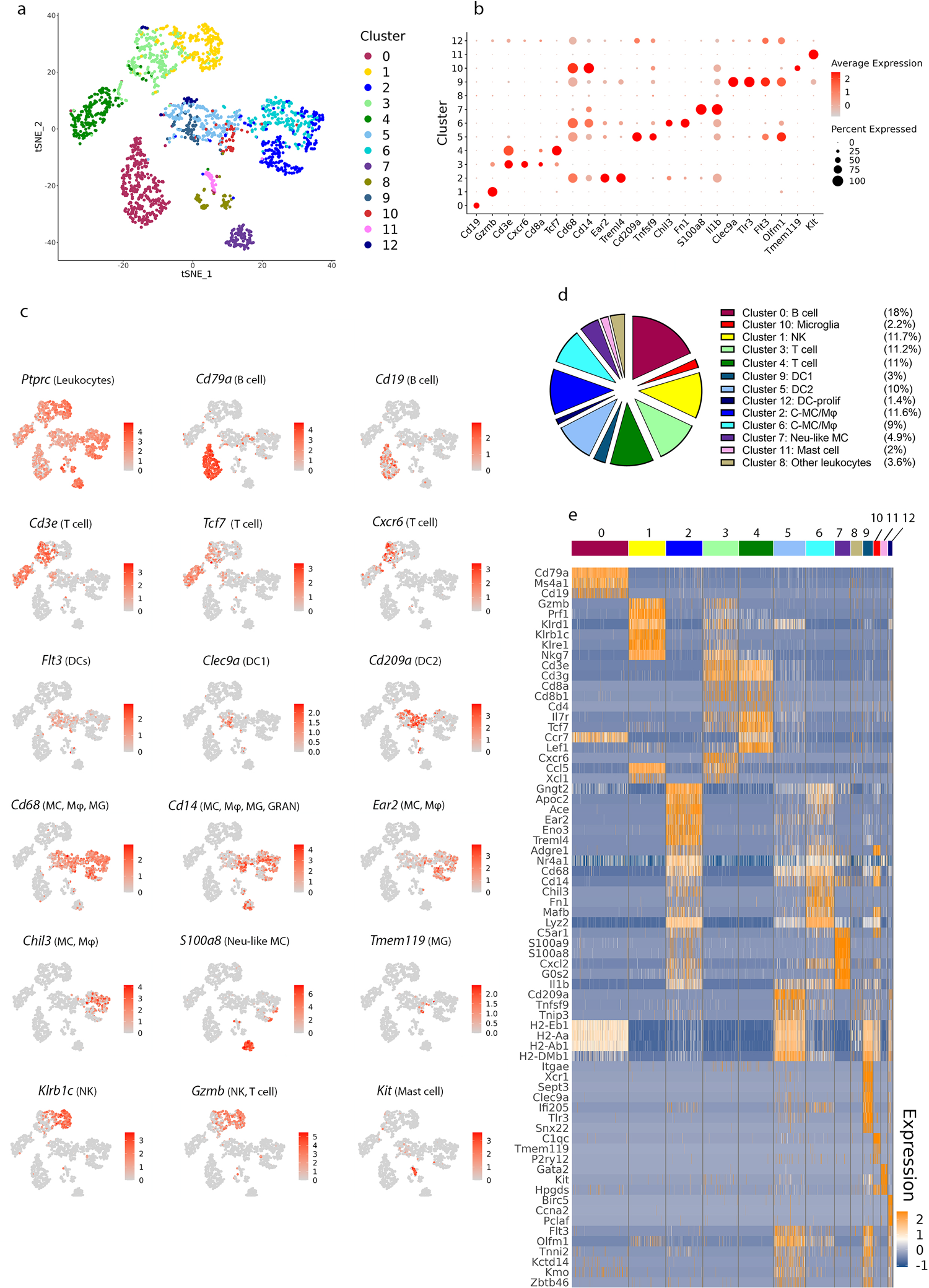
Identifying immune cell types in the mouse choroid based on differentially expressed genes. **(a)** tSNE plot of 2,125 immune cells (CD45^+^) sorted from choroid tissue pooled from n=20 C57Bl/6J mice, showing the thirteen immune cell types that were identified via unsupervised clustering. **(b)** Dot plots showing the expression of cell type-specific genes, with the dot size representing the percentage of cells expressing the gene and the colour representing its average expression within a cluster. **(c)** tSNE maps showing the expression of key genes for the immune cell populations that identified in (b). Grey, low expression; red, high expression, and range of log_2_ normalized counts are shown on the right of each plot. **(d)** Pie charts showing the proportions of different immune cell types within the homeostatic mouse choroid. Clusters were identified based on expression of specific genes (the pie chart legend is ordered according to cell type rather than by numerical order of clusters). The frequency of each cluster is shown as percentage of the total immune cell population to the right of each cluster name. Note that the cluster identified as ‘other leukocytes’ expressed the CD45 gene (*Ptprc*), but none of the other examined lineage markers. **(e)** Heatmap of normalized expression for selected differentially expressed genes in each cluster. Range of log_2_ normalized counts are shown at right hand of heatmap. Blue, low expression; orange, high expression. Monocyte, MC; Macrophages, Mφ; Dendritic cells, DCs; Natural killer cells, NK; Microglia, MG; granulocytes, GRAN; Neutrophil-like monocyte, Neu-like MC.

Three DC clusters were observed in the choroid. Each of these clusters expressed *Flt3* and the DC-specific transcription factor *Zbtb46* (26). The gene expression profiles of these clusters were consistent with conventional type 1 DCs (cDC1, cluster 9), cDC2 (cluster 5), and proliferative DCs (DC-prolif, cluster 12). Both cDC1 and cDC2 clusters expressed antigen processing and presentation related genes such as *H2-Eb1*, *H2-Aa*, *H2-Ab1*, and *H2-DMb1.* cDC1s were distinguished by 302 DEGs including *Itgae*, *Xcr1*, *Sept3*, *Clec9a*, *Pianp, Ffar4*, *Plpp1*, *Ifi205*, *Tlr3*, *Snx22*. The cDC2 cluster was the predominant DC population within the choroid and cells within this cluster were enriched for *Il3ra*, *Cd209a*, *Tnfsf9* and *Tnip3*, and had 252 DEGs. DC-prolif (cluster 12) expressed genes such as *Birc5*, *Ccna2* (27), and *Pclaf* (28) that promote proliferation, in addition to the common DC markers.

Furthermore, two distinct choroidal MC/Mφ) clusters (clusters 2 and 6) and one neutrophil-like monocyte (Neu-like MC) cluster (cluster 7) with unique transcriptomic signatures were resolved in the choroid. Cluster 2 was the predominant MC/Mφ cluster and was distinguished by 689 DEGs including *Gngt2*, *Apoc2*, *Ace*, *Eno3*, and *Treml4*. Choroidal MC/Mφ cluster 6, characterised by 577 DEGs, expressed genes *Chil3*, *Fn1*, *Mafb* and *Lyz2*. Cluster 7 was unique compared to the other choroidal MC/Mφs due to its low expression of *Cd68*, yet enrichment of *Cd14*, *S100a9*, *S100a8*, and *Il1b*. The gene expression of this cluster resembled neutrophil-like monocytes (Neu-like MC), which were recently identified in an anti-glomerular basement membrane crescentic glomerulonephritis mouse model (29, 30); however, further work is needed to determine the identify of this cluster.

Two T cell clusters (cluster 3 and cluster 4; characterised by expression of *Cd3e*, *Cd3g* and *Cd3d*) were identified in the choroid. Cluster 3 expressed genes consistent with CD8^+^ T cells (*Cd8a*, *Cd8b1*, *Nkg7* and *Gzmb*) and was also characterised by DEGs including *Cxcr6* and *Ccl5*. Cluster 4 also expressed *Cd8a*, *Cd8b1* (but not *Nkg7* and *Gzmb*) and *Cd4*. This cluster was enriched for *Il7r* and *Ccr7*, which are expressed by CD8^+^ and CD4^+^ effector memory T cells, as well as *Lef1* and *Tcf7*, which are expressed by naïve T cells and downregulated following TCR engagement. *Tcf7* is a transcription factor that plays an important role during T cell development and differentiation (31, 32). These findings suggest this cluster contains a mixture of T cell subtypes.

### Dendritic cells in the bordering tissues of the mouse brain and retina exhibit tissue-specific transcriptional signatures

Our analysis revealed that the mouse choroid contained three DC clusters, whereas pia mater-enriched tissue contained only one DC cluster. To further characterise DC heterogeneity between these tissues, we compared DEGs between DC clusters. The pia-enriched cDC2 cluster (cluster 4) shared a similar transcriptional signature to the choroidal cDC2 cluster (cluster 5). However, some differences were identified between these two clusters; for example, *Tnfsf9*, *Mt1*, *Tnip3*, and *Ifi30* were highly expressed in choroidal cDC2, while *Cox6a2*, *Bst2*, and *Apoe* was enriched in the pial cDC2 cluster (Figure 4a). Gene ontology (GO) analysis demonstrated that MHC class II genes, *Cd209a*, *Cd74*, *Fcgr2b*, *Lmo1*, and *Ccr2* were involved in the top 30 overrepresented GO terms for both the pial cDC2 cluster and choroidal cDC2 cluster (Figure 4b and d). Cell proliferation, cell adhesion and activation, lymphocyte mediated immunity, antigen processing and presentation, cell differentiation and response to interferon gamma were functions common to pial cDC2 and choroidal cDC2 clusters.

**Figure 4.**
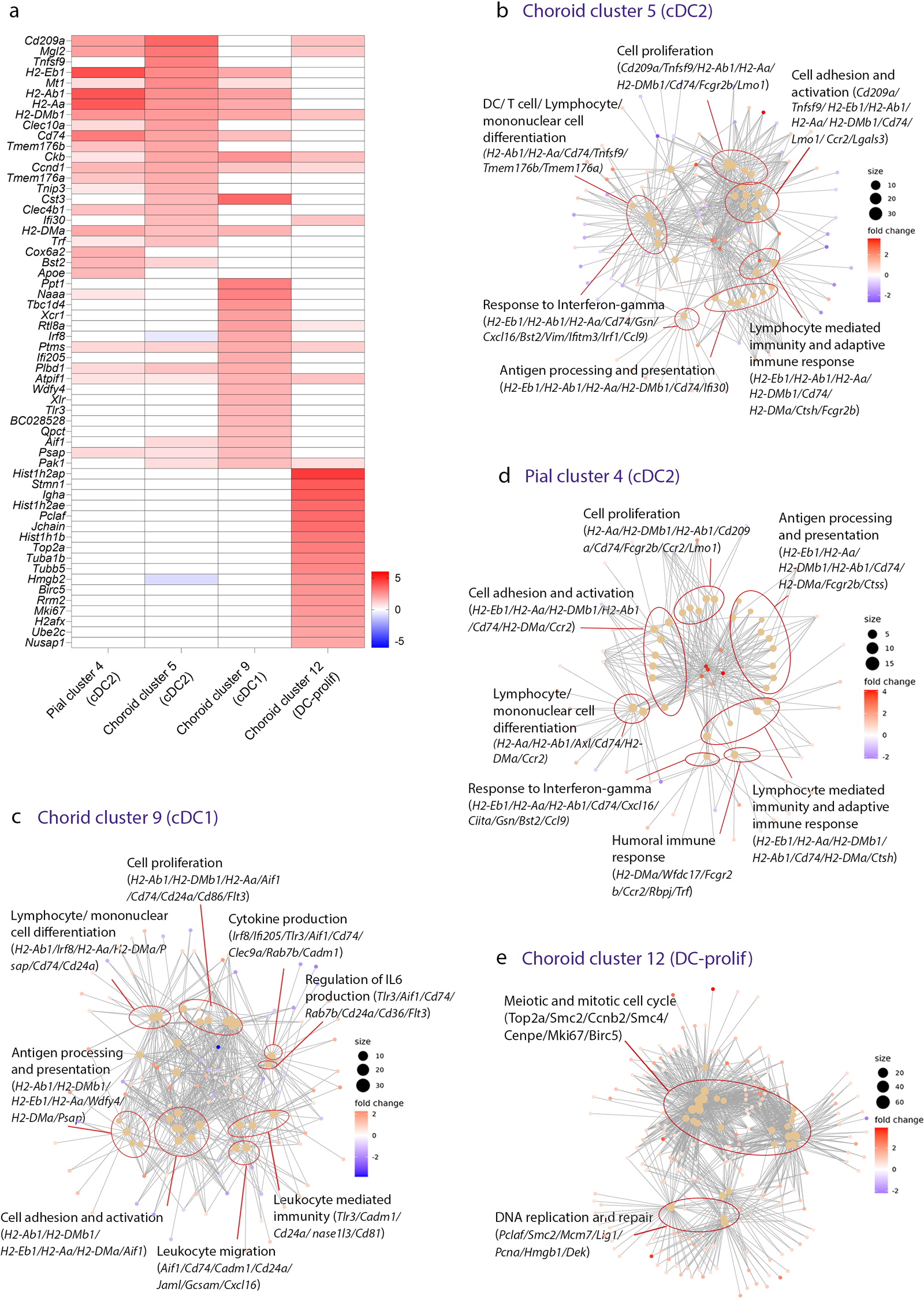
Comparison of overrepresented GO terms of DC clusters within mouse pia mater-enriched tissue and choroid. **(a)** Heat map of normalized gene expression showing the top upregulated DEGs (log2 fold change of >1.5) in pial cDC2 (cluster 4), choroidal cDC2 (cluster 5), and choroidal cDC1 (cluster 9) clusters, and top upregulated DEGs (log2 fold change of >2) in DC-prolif (cluster 12), compared to other clusters in pia mater-enriched tissue and choroid respectively. Range of log_2_ normalized counts are shown to the right of heatmap. **(c-d)** Gene ontology network of top 30 GO terms based on genes that are differentially expressed between each DC cluster and other subsets in either pia mater-enriched tissue or choroid. Each node (filled dots) represents a gene ontology and the size of each node reflects the p values of the terms, with the more significant terms being larger. The main themes within the data are categorized into groups (red circles). A selection of key genes associated with each cluster of gene ontologies is displayed.

Genes involved in the top 30 overrepresented GO terms for the choroidal cDC1 cluster included MHC class II related genes, *Aif1*, *Cd74*, *Cd24a*, *Cd86*, and *Tlr3*; and regulation of IL-6 production was a top 30 GO term that was unique to this cluster (Figure 4c). In contrast, the choroidal DC-prolif cluster (cluster 12) shared few genes with the other characterised DC clusters and was enriched for GO terms related to meiotic and mitotic cell cycle and DNA replication and repair (Figure 4e).

### Mouse pia mater-enriched tissue and choroid contain diverse monocyte/macrophage populations with potential functional differences

To further delineate the heterogeneity of MC/Mφ across these tissues, we conducted a comparison of the top DEGs among MC/Mφ clusters. Striking similarities were observed between choroidal MC/Mφ cluster 2 and pial MC/Mφ cluster 3; and between MC/Mφ cluster 6 and pial MC/Mφ cluster 5. In contrast, the DEGs defining choroidal MC/Mφ cluster 7 and pial MC/Mφ cluster 9 were unique to their respective tissues (Figure 5a).

**Figure 5.**
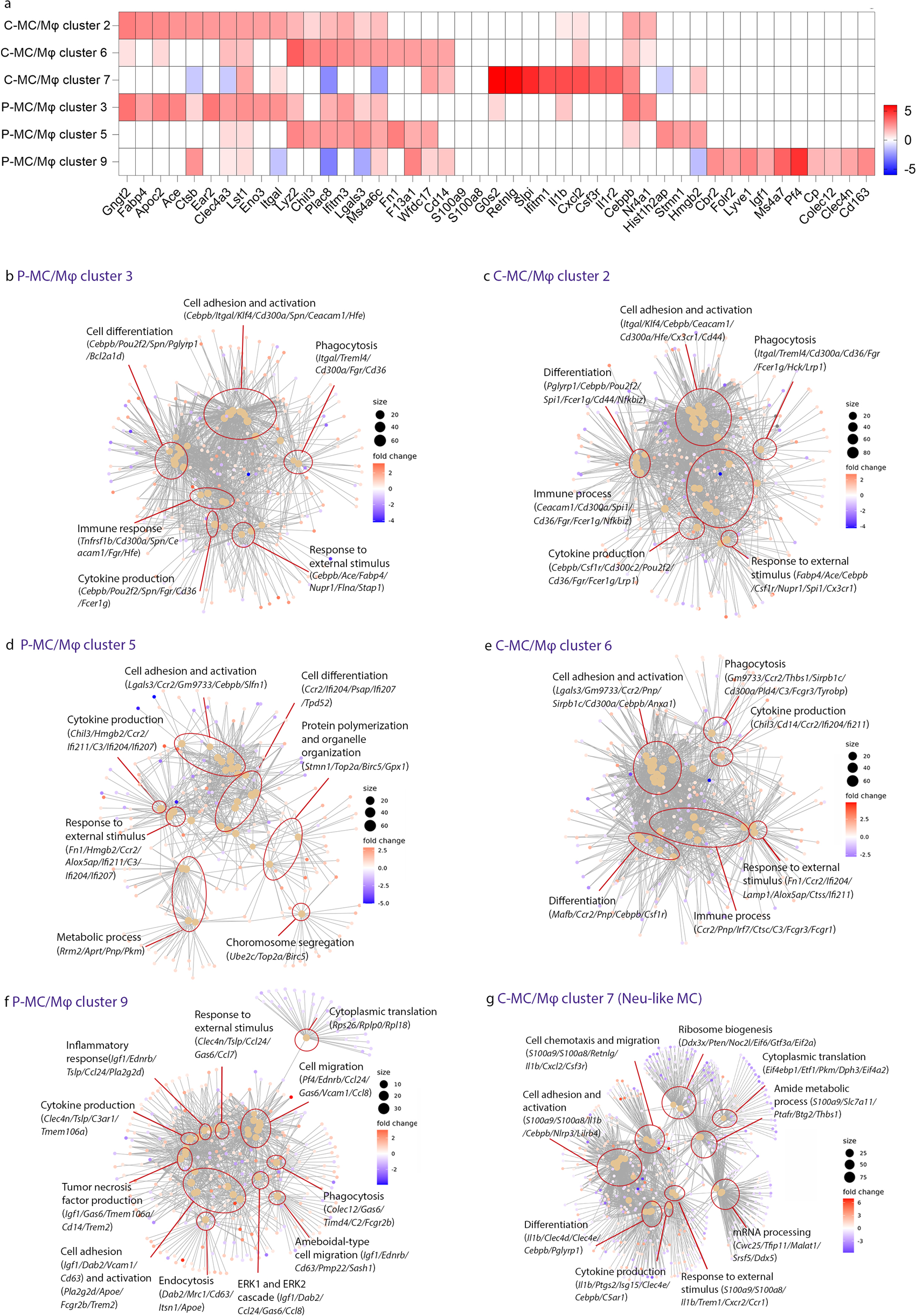
Comparison of overrepresented GO terms of MC/Mφ clusters within mouse pia mater-enriched tissue and choroid. **(a)** Heat map of normalized expression showing the 10-top upregulated DEGs in pia mater-enriched tissue (pial MC/Mφ clusters 3, 5, 9) and choroid (choroidal MC/Mφ clusters 2, 6; Neu-like MC cluster 7), compared to other clusters in the pia mater and choroid respectively. Range of log_2_ normalized counts are shown to the right of heatmap. **(b-g)** Gene ontology networks of the top 30 GO terms based on genes that are differentially expressed by each pial and choroidal MC/Mφ cluster compared to other clusters within the pia mater or choroid respectively. Each node (filled in dots) represents a gene ontology, and the size of the GO terms reflects the p values of the terms, with the more significant terms being larger. The main themes within the data are categorized into groups (red circles). A selection of key genes associated with each cluster of gene ontologies is displayed.

Further comparison of the top overrepresented GO terms revealed common functions between the DEGs of several of the MC/Mφ clusters, including phagocytosis, cell adhesion and activation, cytokine production, response to external stimulus, immune processes and cell differentiation (Figure 5b-g). Pial MC/Mφ cluster 3 and choroidal MC/Mφ cluster 2 showed a high degree of similarity, with the genes *Itgal*, *Klf4*, *Treml4*, *Cd300a*, *Cd36*, *Ace, Fabp4* involved in the top 20 overrepresented GO terms for both clusters (Figure 5b and c). Pial MC/Mφ cluster 5 and choroidal MC/Mφ cluster 6 also displayed similarities, and the top genes involved in the overrepresented GO terms were *Lgals3*, *Gm9733*, *Ccr2*, *Chil3*, *C3*, *Fn1*, *Ifi211* and *Ifi204* (Figure 5d and e). However, pial MC/Mφ cluster 5 uniquely showed an overrepresentation of the GO terms metabolic process, chromosome segregation, and protein polymerization and organelle organization, which suggested an enhanced mitotic state within this cluster (Figure 5d).

The unique pial MC/Mφ cluster (cluster 9) featured *Pf4*, *Dab2*, *Mrc1*, *Gas6*, *Cd63*, *Fn1*, *Ccl24* and *Clec4n* as top genes involved in the overrepresented GO terms (Figure 5f). Unlike other MC/Mφ clusters within pia mater-enriched tissue and choroid, this cluster featured tumour necrosis factor production, endocytosis, amoeboid migration and ERK1/ERK2 cascade among the top overrepresented GO terms. Choroidal MC/Mφ cluster 7, putatively identified as Neu-like MC, showed enrichment of GO terms related to cell chemotaxis and migration, cytokine production, response to external stimulus, mRNA processing, ribosome biogenesis, cytoplasmic translation, and metabolic processes. *S100a9*, *S100a8*, *Il1b*, *C5ar*, *Cxcl2*, *Cxcr2*, *Csf3r* were the top genes related to these overrepresented pathways (Figure 5g).

### Heterogeneous B and T cell gene expression profiles within mouse pia mater-enriched tissue and choroid

B cells constituted a sizeable proportion of immune cells detected within the healthy mouse choroid and pia mater-enriched tissue. Although mice were perfused to clear the vasculature of blood components, including circulating leukocytes, prior to tissue collection complete removal of peripheral components cannot be guaranteed (33). Hence, there remains a possibility that a portion of lymphocytes detected in this study may represent contaminating peripheral immune cells. Regardless, immunostaining and confocal microscopy demonstrated the presence of CD19^+^ B cells *in situ* within the homeostatic mouse pia mater and choroid (Figure 6a).

**Figure 6.**
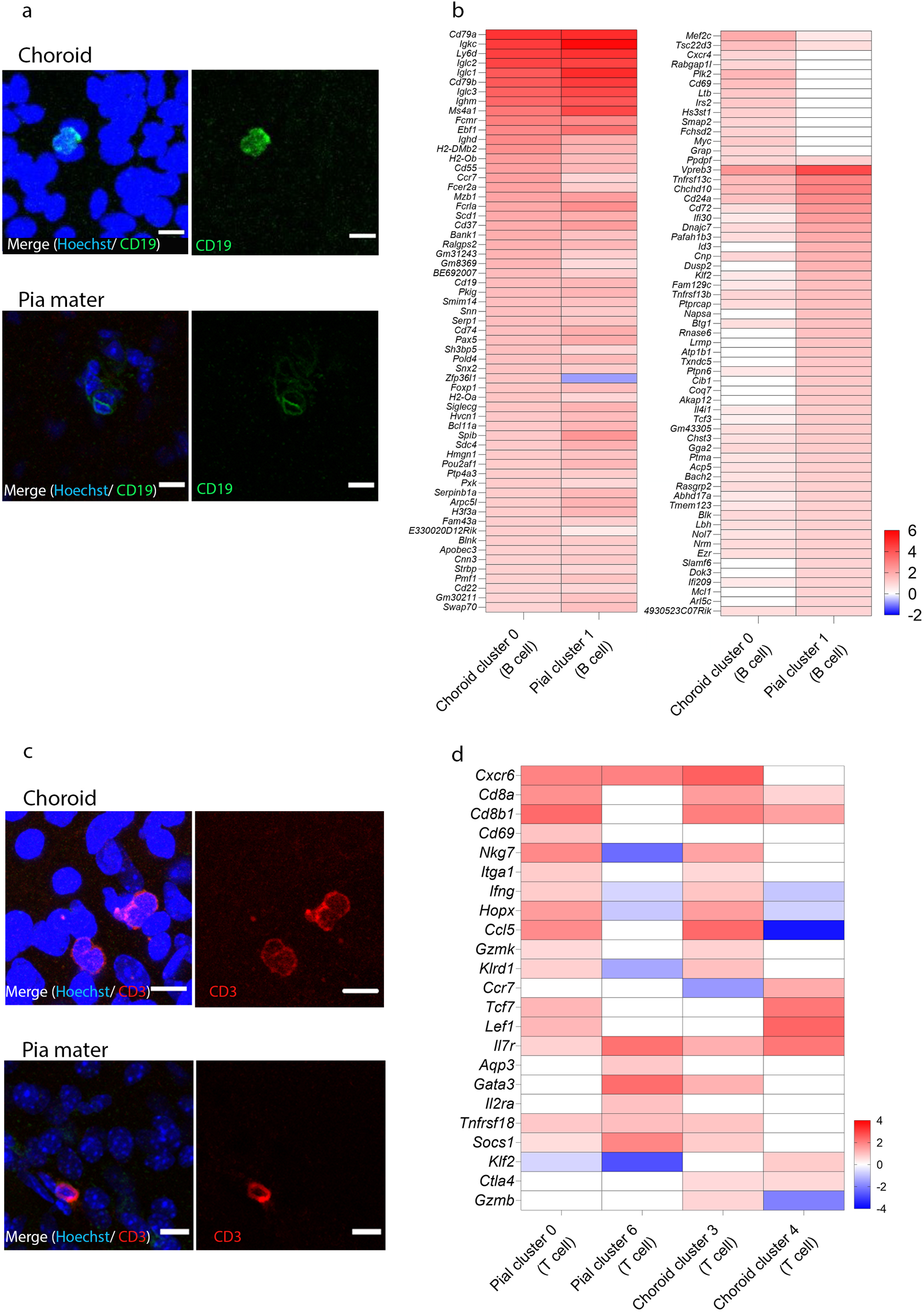
Comparison of lymphocyte cluster DEGs within mouse pia mater-enriched tissue and choroid. **(a)** Identification of B cells within the healthy mouse (n=1) choroid and pia mater using fluorescent immunohistochemistry and confocal microscopy. Immunostaining panels display expression of CD19 (green), and blue nuclear staining (Hoechst). Scale bars represent 10 µm**. (b)** Heatmap of normalized expression for top upregulated DEGs (log_2_ fold change of >1.0) in the choroid B cell (cluster 0) compared to other clusters in the choroid; and pial B cell (cluster 1) compared to other clusters in the pia mater. Range of log_2_ normalized counts are shown to the right of heatmap. **(c-d)** Identification of T cells within the healthy mouse choroid and pia using fluorescent immunohistochemistry and confocal microscopy. Immunostaining panels display expression of CD3 (red), and blue nuclear staining (Hoechst). Scale bars represent 10 µm**. (d)** Heatmap of normalized expression of T cell genes pia (cluster 0 and 6) compared to other clusters in the pia; and in the choroid (cluster 3 and 4) compared to other clusters in the choroid. Range of log_2_ normalized counts are shown to the right of heatmap.

The choroidal B cell cluster (cluster 0) was distinguished by 819 DEGs compared to other choroidal clusters, whereas the pial B cell cluster (cluster 1) had 771 DEGs relative to other pial clusters. The top upregulated DEGs (log_2_ fold change of >1.0) were compared between the pia mater and choroidal B cell clusters, and it was found that pan-B cell markers including *Cd79a*, *Cd79b* and *Cd19* and other B cell markers such as *Igkc*, *Iglc2*, *Iglc2*, *Iglc3* and *Ighm* were upregulated in B cell clusters in both tissues. Mature B cell markers such as *Ms4a1*, *Fcer2a*, and *Bank1* were also upregulated in both B cell clusters. However, there were some tissue-specific DEGs that were upregulated in choroidal and pial B cell clusters respectively. For example, *Cxcr4*, *Rabgap1l*, *Plk2*, *Cd69*, *Ltb*, *Irs2*, *Hs3st1*, *Smap2*, *Fchsd2*, *Myc*, and *Grap* were exclusively upregulated in the choroidal B cell cluster. While *Id3*, *Dusp2*, *Napsa*, *Rnase6*, *Lrmp*, *Atp1b1*, *Txndc5*, *Cib1*, *Coq7*, *Akap12*, *Slamf6*, *Dok3*, *Mcl1*, and *Arl5c* were DEGs unique to the pial B cell cluster (Figure 6b). These differentially expressed genes between pial and choroidal B cell clusters are involved in GO terms including the cellular process, localization, response to stimulus, signalling, developmental process, metabolic process, and immune system process.

Unsupervised clustering of scRNA-seq data revealed that the mouse choroid and pia mater-enriched tissue contained various T cell clusters (Figures 2 and 3). Similar to B cells, immunostaining and confocal microscopy confirmed that the naïve mouse choroid and pia mater contained CD3^+^ T cells within the tissue (Figure 6c). To characterise T cell heterogeneity between these tissues, we compared top upregulated DEGs between pial T cell clusters (clusters 0 and 6) and choroidal T cell clusters (clusters 3 and 4) (Figure 6d). Pial T cell cluster 0 shared a similar DEGs to the choroidal T cell cluster 3, with *Cd8a*, *Cd8b1*, *Nkg7, Il7r, Cxcr6*, *Itga1*, *Ifng*, *Hopx*, *Gzmk*, *Klrd1*, and *Ccl5* among the top upregulated DEGs for both clusters. However, some tissue-specific differences were observed, for example, *Cd69*, *Tcf7* and *Lef1* were upregulated by pial T cell cluster 0 (but not choroidal T cell cluster 3), whereas *Ctla4 and Gzmb* were enriched in the choroidal T cell cluster 3 (but not the pial T cell cluster 0). Interestingly, the pial T cell cluster 6 had a distinct transcriptional profile from the pial cluster 0 and both choroidal T cells clusters, and was enriched for *Aqp3*, *Socs1*, *Il2ra* and *Gata3*.

### Transcriptomic profile of leukocytes within the human choroid

Having established a transcriptomic profile of immune cells within the mouse choroid and pia mater-enriched tissue, we next sought to determine if similar populations were present in human tissues. We therefore performed scRNA-seq on immune cells (CD45^+^) isolated from the choroid of a 45-year-old male eye donor. Unsupervised clustering and tSNE projections were performed on 6,501 cells sequenced from the human choroid (70,322 mean reads per cell) (Figure 7a). Clusters were identified based on cell-specific gene expression profiles. Similar to the mouse choroid, the human choroid contained various immune cell types including B cells, T cells, DC, MC/Mφ, NK cells, and mast cells (Figure 7b-d).

**Figure 7.**
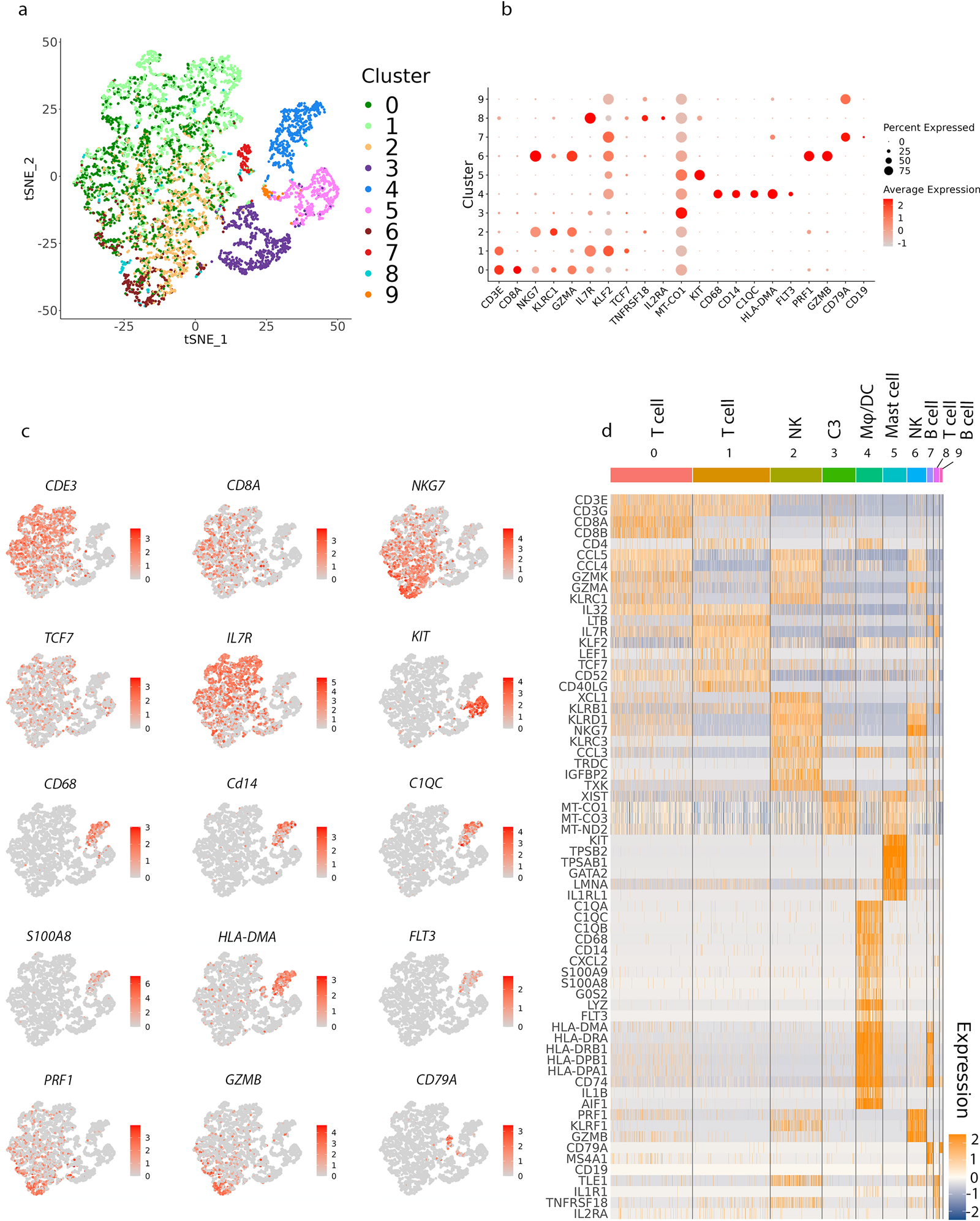
Identifying immune cell types in the human choroid based on differentially expressed genes. **(a)** tSNE plot of 6,501 immune cells (Cd45+) (n=1 donor) showing the ten immune cell types that were identified via unsupervised clustering. **(b)** Dot plots showing the expression of cell type-specific genes, with the dot size representing the percentage of cells expressing the gene and the colour representing its average expression within a cluster. **(c)** tSNE maps showing the expression of key marker genes for the immune cell populations that identified in (b). Grey, low expression; red, high expression. The range of log2 normalized counts are shown to the right of each plot. **(d)** Heatmap of normalized expression for selected genes in each cluster. Range of log_2_ normalized counts are shown to the right of heatmap. Blue, low expression; orange, high expression.

T cells were the predominant immune cell cluster identified in the human choroid. However, this may not accurately represent the composition of immune cells *in situ*, as (unlike the mouse choroid) it was not possible to exclude circulating leukocytes from the human choroid prior to tissue processing. Three T cell clusters (cluster 0, cluster 1 and cluster 8) were identified in the human choroid. Cluster 8 was a small cluster that expressed *IL7R, KLRB1, TXK, IL1R1, TNFRSF18, TLE1, and IL2RA.* T cell cluster 0 expressed genes consistent with cytotoxic CD8^+^ T cells (*CD8A*, *CD8B1*, *NKG7*, *GZMA, and KLRC1*), whereas T cell cluster 1 expressed genes consistent with naïve T cells and CD4^+^ effector memory T cells, such as IL*7R*, *KLF2*, *TCF7*, and *LEF1* and *CD4*. Whilst several MC/Mφ and DC clusters were identified in the mouse choroid, only one cluster (cluster 4) expressed genes characteristic of MC/Mφ (*CD14, CD68*) and DCs (*FLT3*) in the human choroid (Figure 7b-d). This cluster was also characterised by expression of *HLA-DMA*, *HLA-DRA*, *HLA-DRB1*, *HLA-DPB1*, *HLA-DPA1*, and *CD74,* suggesting that cells within this cluster likely function in antigen processing and presentation. It is possible that with increased sequencing depth, this cluster may resolve into individual MC/Mφ and DC populations as observed in the mouse choroid. In addition, *S100A9* and *S100A8* were upregulated in a part of this cluster. Cluster 3 (C3) had high level of mitochondrial gene expression such as *MT-CO1* and *MT-CO2* which could be a result of high fraction of apoptotic or lysing cells in the sample given the delay between donor time of death and sample processing.

Overall, these findings demonstrated that the human choroid contained MC/Mφ/DCs, NK cells, mast cells, T cells and B cells (Figure 7d). However, due to differences in cell numbers sequenced, sequencing depth and inability to perfuse human tissue prior to processing, direct comparison of clusters and their transcriptomic signatures was not possible between human and mouse tissues.

## DISCUSSION

This study used scRNA-seq to generate a transcriptomic cell profile of immune cells within the healthy mouse pia mater-enriched leptomeninges and choroid. Multiple immune cell types were identified in these homologous CNS bordering tissues, with some clusters common to both tissues and others exhibiting tissue-specific gene expression signatures. Myeloid cells were the predominant immune cells detected within mouse pia mater-enriched tissue and choroid, consistent with previous literature demonstrating populations of macrophages and DCs within these tissues (9, 10, 34). An important outcome of this study is the identification of unique sets of genes for each of the identified pial and choroidal macrophage and DC clusters, which may be useful for defining cell-specific markers for these subtypes. This may enable future studies to map the spatial distribution of macrophage and DC subtypes and expand our understanding of their role in healthy conditions and during inflammatory diseases of the CNS.

Of the three MC/Mφ clusters identified in pia mater-enriched tissue, pial MC/Mφ cluster 9 expressed genes consistent with *Cd163^+^* Mφs, which were previously reported in the subdural meninges (9). Interestingly, this cluster showed DEGs related to amoeboid migration, which was unique among pial macrophage clusters. The mobility of immune cells within the meningeal spaces, such as T cells and antigen presenting cells, and their migration to other sites has been demonstrated previously (35–38). Our data potentially extend these findings to include pial macrophages, although further investigation is warranted. The choroid also contained three MC/Mφ clusters but in contrast to pia mater-enriched tissue, the choroid did not contain a cluster that resembled *Cd163^+^*Mφs. The differences in gene expression observed between pial and choroidal MC/Mφ clusters is consistent with previous studies demonstrating differences in the ontogenies of macrophages within the subdural meninges and choroid. Whilst subdural meningeal macrophages are prenatally-derived, long-lived cells that are maintained through *in situ* self-renewal (8), choroidal macrophages are short-lived monocyte-derived cells that are replenished from the bone marrow (39).

However, we did identify some similarities between selected myeloid clusters in pia mater-enriched tissue and choroid. Pial MC/Mφ cluster 3 and choroidal MC/Mφ cluster 2 appeared to be equivalent macrophage populations, sharing common top upregulated DEGs and overrepresented GO terms. Additionally, both the pia mater-enriched tissue and choroid contained a cDC2 cluster with similar transcriptomic signatures. The choroid also contained cDC1, DC-Prolif and Neu-like MC clusters, which were absent in pia mater-enriched tissue. It is possible that a greater number of myeloid clusters within the choroid may be required for enhanced surveillance due to the different vascular properties of this tissue (fenestrated blood vessels) compared to the pia mater (non-fenestrated blood vessels) (4, 40). A limitation of our study though, is that a larger number of cells were sequenced from the choroid (2,125 cells) compared to pia mater-enriched tissue (1,418 cells). It is therefore possible that the reduced number of cell types identified in pia mater-enriched tissue compared to choroid is due to the lower number of immune cells sequenced. Despite this, an advantage of our study is the high sequencing depth achieved for mouse pia mater-enriched tissue and choroid, which enables higher accuracy in determining the true transcriptional profile of cells (41). Another unique feature of our scRNA-seq study was that our tissue dissection and cell sorting strategy limited the number of non-target microglia cells within the datasets, as it was our intention to characterise immune populations within the bordering tissues of the CNS, not the microglia that reside within the neural parenchyma.

In pia mater-enriched tissue we identified a distinct cluster of neutrophils. A previous study reported that the dura mater and pia mater contained a large population of neutrophils, and that most of these cells were *bona fide* extra-vascular cells and not contaminants from the circulation or lumen of blood vessels (42). Although the role of neutrophils in neuroinflammation has been investigated (43), their function in the healthy meninges is not well understood. It is hypothesised that neutrophils residing within the tissues need to be regulated to avoid degranulation and prevent CNS damage (44). Whilst we did identify a putative Neu-like MC cluster in the mouse choroid, classical neutrophils were exclusively identified in pia mater-enriched tissue and not the choroid. Neu-like MC are granulocyte monocyte progenitor (GMP)-derived neutrophil-like monocytes that produce proinflammatory cytokines such as IL-1β (29, 30). Recently, Chen *et al.* found a unique Neu-like MC cluster, similar to that described in our study, in an anti-glomerular basement membrane crescentic glomerulonephritis mouse model (30). To the best of our knowledge, this is the first study to identify a putative Neu-like MC population in the choroid. Given that Neu-like MCs play a role in disease in other organ systems, future studies of choroidal Neu-like MCs may reveal a novel role for these cells in ocular inflammation, although further work is required to validate the identity and function of these cells.

We identified B cell, T cell and NK cells clusters within both pia-enriched tissue and the choroid using scRNA-seq and demonstrated the presence of B cell and T cells in these tissues using immunohistochemistry. These findings are consistent with a previous scRNA-seq study that showed B cell, T cell, and NK cell populations within the healthy mouse dura mater, enriched subdural meninges and choroid plexus (9). T cells were also identified in the retinal pigment epithelium/choroid using single cell RNA sequencing in the mouse (45) and human (46). When compared to previous studies of the subdural meninges (9), our data showed a higher proportion of B and T cells within pia mater-enriched tissue. These differences may be explained by differences in experimental conditions, for example - such as different mouse sexes used between these two studies or differences in animal housing environments. However, we also cannot exclude the possibility that a portion of lymphocytes identified in our study may be peripheral contaminants, as previously mentioned. Future studies assessing the origin and density of lymphocyte populations within these tissues are required to validate the *in situ* distribution of B and T cell clusters.

Within the CNS and its bordering tissues, most studies have focused on microglia, macrophages, and T cells, whereas little is known about B cells. Recent scRNA-seq studies have substantiated the concept of long-term residence and local development of B cells within the meninges (33, 47, 48). Specifically, these studies have revealed that meningeal B cells derive from calvarial bone marrow and primarily congregate near the dural sinuses, where endothelial cells express crucial niche factors conducive to B cell development (48). Here, we demonstrated that B cells are also present within the mouse pia mater-enriched tissue and choroid, as well as the human choroid. However, it is important to highlight that in our study, the pial and choroidal B cells expressed pan mature B cell markers but not genes associated with B cell development. Whether pial B cells are derived from the calvarial-meningeal pathway of B cell development (47), or from systemic circulation remains to be determined.

Comparison of B cell clusters within the healthy mouse pia mater and choroid revealed that *Cxcr4* is highly expressed by the choroidal B cell cluster but not the pial B cell cluster. *Cxcr4* plays a crucial role in regulating the homeostasis of B cells and humoral immunity, as inactivation of Cxcr4 has been linked to decreased numbers of B cells and defective T cell-independent responses in the peritoneal cavity (49). Additionally, the choroidal B cell cluster differentially expressed *Cd69*, a molecule that triggers B cell activation and the maturation of DCs, via direct cellular contact, that have an amplified capacity to promote Th2 responses (50). Genes that were enriched in the pial B cell cluster (but not expressed by the choroidal B cell cluster) included *Id3*, *Dok3* and *Slamf6*. *Id3*, a transient inhibitor of E protein activity (51), is known to be expressed by naïve B cells and plays a role in the ability of activated B cells to undergo expansion, class switching recombination and germinal centre development (52). Upon exposure to antigens, downregulation of *Id3* results in the release of E2A and E2-2 proteins that are essential for antigen-induced B cell differentiation (53). *Dok3* is an adapter molecule involved in the recruitment of inhibitory molecules, which is suggested to play a role in negative regulation of immunoreceptor signalling in B cells and macrophages (54). Finally, *Slamf6* is involved in the interaction between naïve B and T cells. This interaction results in upregulation of T cell cytokine migration inhibitory factor, leading to augmented expression of its receptor CD74 on B cells, thereby enhancing B cell survival (55). The variations in gene expression between pial and choroidal B cells suggest nuanced molecular signatures underlining the specialized functions of B cell clusters in distinct anatomical locations within the CNS supporting tissues.

We also performed scRNA-seq of immune cells isolated from human choroid-RPE. Similar to the mouse choroid, the human choroid contained DC, macrophages, NK cells, mast cells, T cells and B cells. These findings highlight the diverse immunological landscape of the human choroid; however, the interpretation of these data are limited due to cells being obtained from only n=1 donor, a lack of perfusion to remove circulating leukocytes from the vasculature of the human choroid, and the time delay between tissue retrieval by tissue bank personnel and processing for scRNA-seq (17 hours).

In conclusion, scRNA-seq revealed that mouse pia mater-enriched tissue and choroid contained similar immune cell types; however, tissue-specific immune cell clusters and gene expression signatures were also characterised. These differences may be explained by the different vascular properties of the choroid (leaky fenestrated blood vessels) compared to the pia (non-fenestrated blood vessels with tight junctions) (4, 40). However, recent studies demonstrating a role for immune cells in maintaining tissue homeostasis may indicate that the differences in immune environments between the pia mater and choroid represent functional specialisations to support the underlying brain and retina respectively. Our study has several limitations, therefore further research is required to validate the identified immune cell clusters, their spatial distribution and function. Overall, our findings serve as a base for understanding the diversity of resident immune cells in the CNS bordering tissues.

## DECLARATIONS

### Ethics approval and consent to participate

Mouse experiments were approved and performed in accordance with the QIMR Berghofer Medical Research Institute Animal Research Ethics Committee (approval A18613M); QUT AEC administrative approval 1800001261. A notifiable low risk dealing for the use of genetically modified mice in this project was obtained from the QUT University Biosafety Committee (NLRD approval 1800000957). Human tissue experiments were approved by Metro South Human Research Ethics Committee (HREC/07/QPAH/048); QUT HREC administrative approval 0800000807/ERM Project No. 5297.

### Consent for publication

Not applicable. De-identified human tissue was obtained for scRNA-seq studies.

### Competing interests

The authors declare that they have no competing interests.

### Funding

This work was funded by an Australian Research Council Discovery Early Career Research Award (DE180101075) to S.D., and a QUT inter-program collaborative grant to S.D. and D.G.H.

### Authors’ contributions

Conceptualization, F.E, S.J.D.; Methodology, F.E., P.W., D.G.H, S.J.D.; Investigation, F.E., P.W., D.G.H., S.J.D.; Resources, S.J.D.; Writing – original draft, F.E., P.W., S.J.D.; Writing – review & editing, F.E., P.W., D.G.H, S.J.D.; Supervision, S.J.D., D.G.H.; Funding acquisition, S.J.D., D.G.H.

## Acknowledgements

The authors thank the facilities, scientific and technical assistance of QUT Central Analytical Research Facility, QUT eResearch and QIMR Berghofer Animal Facility.

## Key resources table

| REAGENT<br>or<br>RESOURCE | SOURCE | IDENTIFIER | Concentration | Working<br>concentration/dilution |
| --- | --- | --- | --- | --- |
| Antibodies |  |  |  |  |
| Rat anti-mouse<br>CD45 BV421 | BD<br>Biosciences | 563890 | 0.2 mg/mL | 0.25 µg per million cells |
| Purified rat anti-mouse CD16/CD32<br>(Fc Block) | BD<br>Biosciences | 553142 | 0.5 mg/mL | 0.5 µg per million cells |
| Mouse Anti-Human<br>CD45 antibody<br>BV421 | BD<br>Biosciences | 563879 | - | 2.5 µL per million cells |
| Human Fc block | BD<br>Biosciences | 564220 | 0.5 mg/mL | 2.5 µg per million cells |
| CD19 polyclonal<br>antibody | Invitrogen | PA5-114970 | 1 mg/ml | 1/200 |
| CD3 monoclonal<br>antibody | Invitrogen | 14-0032-82 | 0.5 mg/mL | 1/400 |
| Hoechst 33342<br>Solution (20 mM) | Life<br>Technologies | 62249 | 20 mM | 1/2000 |
| Goat anti-rabbit IgG<br>(H+L) Alexa Fluor<br>594 | Life<br>Technologies | A11012 | 2 mg/mL | 1/400 |
| Chemicals |  |  |  |  |
| Paraformaldehyde<br>(PFA) | Sigma Aldrich | P6148-500G |  |  |
| Propidium iodide | BD<br>Biosciences | 556463 | 50 µg/mL |  |
| Collagenase type II | Life<br>Technologies | 17101-015 |  |  |
| HBSS (calcium, magnesium, no phenol red) | Life Technologies | 14025092 |  |  |
| DMEM/F12 | Gibco | 11320033 |  |  |
| Bovine serum albumin | Sigma Aldrich | A7906-500G |  |  |
| DMSO | Sigma Aldrich | D2438-5x10ML |  |  |

---

**Supplementary Table 1.**
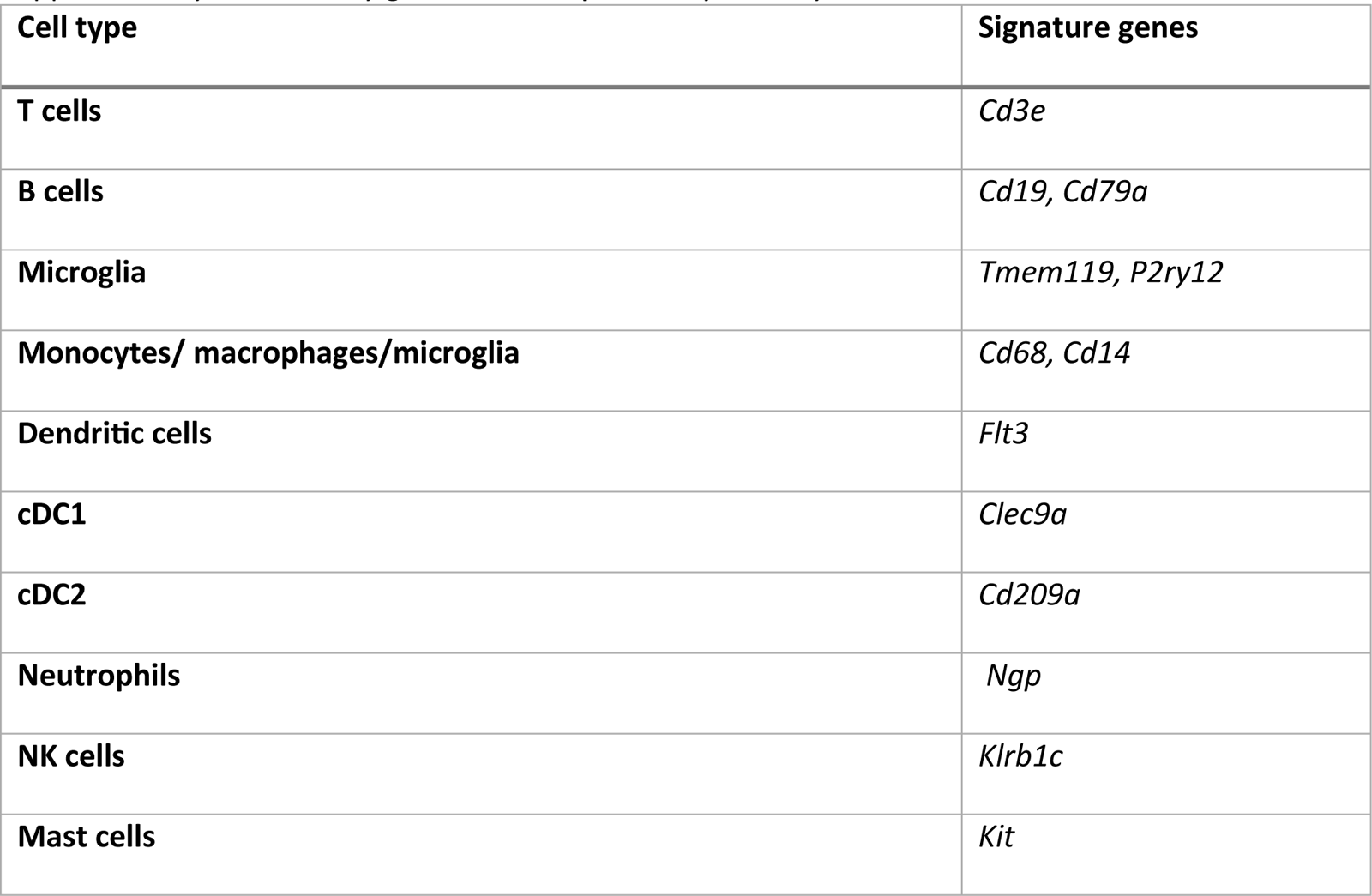
Key genes used to putatively identify immune cell clusters.

| Cell type | Signature genes |
| --- | --- |
| T cells | *Cd3e* |
| B cells | *Cd19, Cd79a* |
| Microglia | *Tmem119, P2ry12* |
| Monocytes/ macrophages/microglia | *Cd68, Cd14* |
| Dendritic cells | *Flt3* |
| cDC1 | *Clec9a* |
| cDC2 | *Cd209a* |
| Neutrophils | *Ngp* |
| NK cells | *Klrb1c* |
| Mast cells | *Kit* |

## Notes

https://www.ncbi.nlm.nih.gov/geo/query/acc.cgi?acc=GSE253419

### Competing Interest Statement

The authors have declared no competing interest.

